# The diversity and distribution of endophytes across biomes, plant phylogeny, and host tissues—how far have we come and where do we go from here?

**DOI:** 10.1101/793471

**Authors:** Joshua G. Harrison, Eric A. Griffin

**Affiliations:** Department of Botany, University of Wyoming, Laramie, WY 82071, USA; New Mexico Highlands University, Las Vegas, NM 87701, USA

**Keywords:** endophytes, fungal endophytes, bacterial endophytes, phyllosphere, microbial ecology, biodiversity, biogeography, plant-microbe interactions

## Abstract

The interiors of plants are colonized by diverse microorganisms that are referred to as endophytes. Endophytes have received much attention over the past few decades, yet many questions remain unanswered regarding patterns in their biodiversity at local to global scales. To characterize research effort to date, we synthesized results from ∼600 published studies. Our survey revealed a global research interest and highlighted several gaps in knowledge. For instance, of the seventeen biomes encompassed by our survey, seven were understudied and together composed only 7% of the studies that we considered. We found that fungal endophyte diversity has been characterized in at least one host from 30% of embryophyte families, while bacterial endophytes have been surveyed in hosts from only 10.5% of families. We complimented our survey with a vote counting procedure to determine endophyte richness patterns among plant tissue types. We found that variation in endophyte assemblages in above-ground tissues varied with host growth habit. Stems were the richest tissue in woody plants, whereas roots were the richest tissue in graminoids. For forbs, we found no consistent differences in relative tissue richness among studies. We propose future directions to fill the gaps in knowledge we uncovered and inspire further research.

**Originality-Significance Statement:** Much remains to be learned regarding the biodiversity and distribution of the microbes that colonize the interiors of plants. Here, we surveyed approximately 600 publications to characterize gaps in knowledge and provide a roadmap for future research. We compared biomes, plant families, and geographical regions in terms of the research interest that they have garnered. Additionally, we synthesized published results and report that variation in endophyte richness among plant tissue types is a function of host growth habit. Stems were the richest tissue in woody plants, whereas roots were the richest tissue in graminoids. We hope to inspire research to fill the gaps in knowledge that we uncovered.

## Introduction

In 1887, Galippe reported that soil-derived microbes could reside within the above-ground tissues of healthy plants. At the time, this work was underappreciated, perhaps because of the long-prevailing attitude that microbial assemblages solely comprised deleterious pathogens (Compant et al. 2012), this notwithstanding the work of A. B. Frank and others demonstrating the mutualistic nature of the mycorrhizal-host relationship (Frank 1885; Frank 2005; Trappe 2005). Nevertheless, Galippe’s observations set the stage for an exploration of the non-pathogenic portion of the plant microbiome that took place from the late 1800s into the mid 1900s (Auret 1930; Hardoim et al. 2015; Janse 1897; Laurent 1889; Rayner 1915). During those decades, knowledge began to accumulate regarding the diversity, prevalence, and ecological roles of so called “endophytes” (Box 1; Campbell 1908; Hyde and Soytong 2008), with much early work focused on the fungi living within grasses (e.g., Neill 1940; Sampson 1937). Seminal research in the 1970s and 80s led to widespread acknowledgement of the ubiquitous nature of non-pathogenic fungi and bacteria in plant tissues, particularly within leaves (Carroll and Carroll 1978; Carroll 1988; Petrini 1991). These studies have inspired an ever-growing interest from microbial ecologists (Fig. 1), yet answers to many basic questions regarding the natural history, biogeography, ecology, and evolution of endophytes remain elusive.

**Figure 1:**
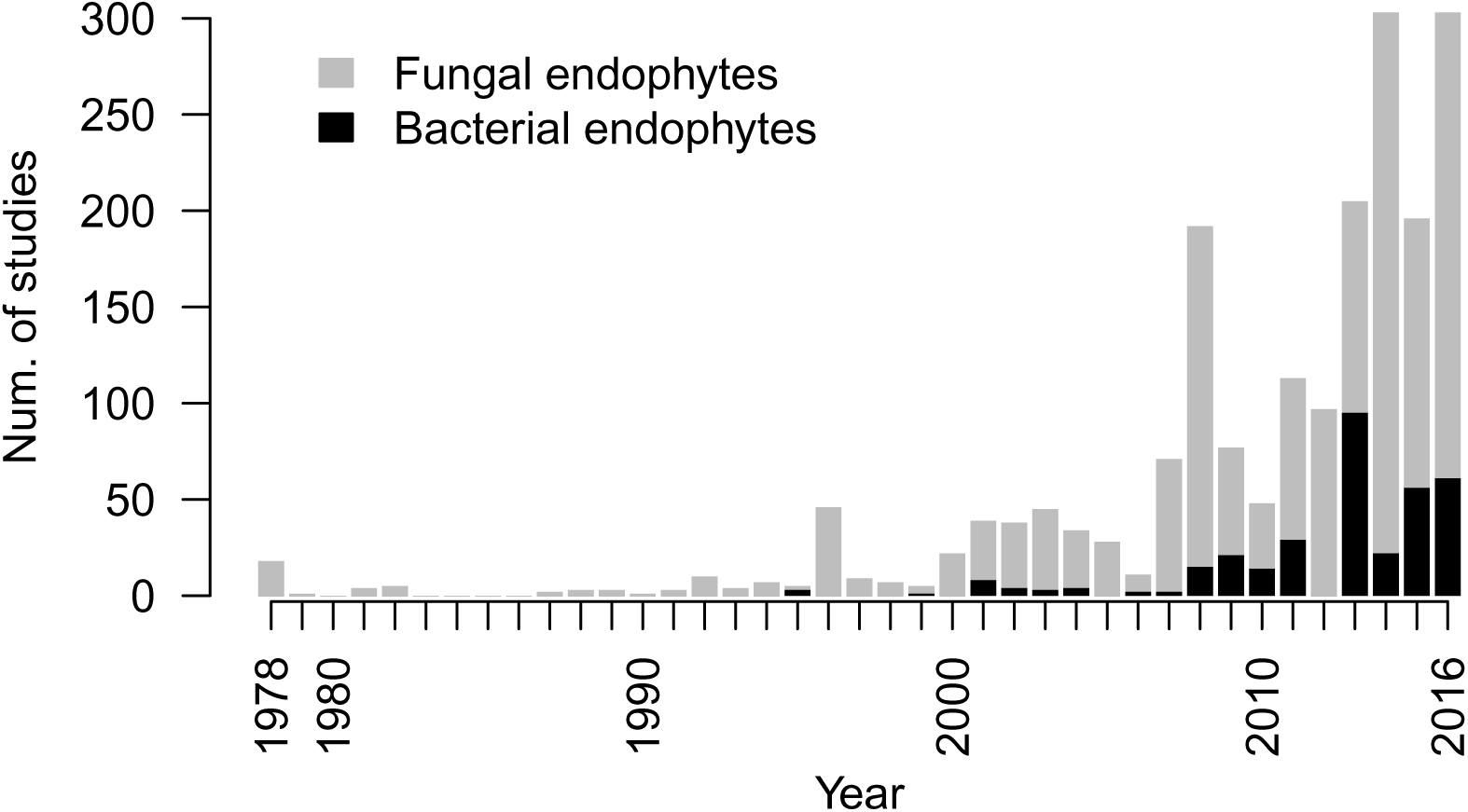
The number of studies characterizing endophyte biodiversity published each year since the late 1970s. Studies are parsed by taxonomy with fungal studies in gray and bacterial studies in black.

#### Box 1. What, exactly, is an endophyte?

The term ‘endophyte’ is believed to have originated with de Bary (1866), who so dubbed pathogenic, plant-inhabiting microbes, because of their habitat (also see Link 1809). Since then, the term endophyte has been expanded to invoke both a habitat and a non-pathogenic lifestyle (at least in some hosts and life history stages), and encompasses fungal (Rodriguez et al. 2009; Petrini 1991), bacterial (Griffin and Carson 2015; Ryan et al. 2008), and archael taxa (Moissl-Eichinger et al. 2018; Müller et al. 2015). The term endophyte can be used *sensu lato* to refer to those taxa that live inside of plant tissues, either inside of or between host cells. However, in our experience, contemporary microbial ecologists most often use the term *sensu stricto* to refer to those taxa which over some portion of their life history, do not cause obvious harm to their hosts, such as inducing a hypersensitive response (Wilson 1995; Petrini 1991; Stone et al. 2000). The lack of precision in this definition is somewhat unsatisfying, but does hint at the complex life histories of many endophytic taxa (Rodriguez et al. 2009). Indeed, for perhaps the majority of endophytic taxa, individuals are horizontally transmitted among hosts and, consequently, may exist outside of the plant corpus for some time, for instance as spores or endospores, free living cells or colonies, or as epiphytic fruiting bodies on decaying tissue (Malloch and Blackwell 1992; Rodriguez et al. 2009). The term endophyte is particularly strained by the mycorrhizal fungi, which possess a mycelium that grows externally to the host but that also penetrates the root epidermis (Schulz and Boyle 2006; Jumpponen 2001). The categorization of these fungi as endophytes seems to be on an author-by-author basis (Schulz and Boyle 2006). These examples illustrate how the term endophyte is useful for communication, but not biologically well-delineated.

What is clear, however, is that fungal and bacterial endophytes are important—even critical—components of the world’s ecosystems. Endophytes can affect plant phenotype, including decreasing disease susceptibility (Arnold et al. 2003; Busby et al. 2016; Christian et al. 2017; Compant et al. 2005; Herre et al. 2007), increasing resistance to abiotic stressors (Márquez et al. 2007; Redman et al. 2002; Rodriguez et al. 2008), shaping phytochemical profiles (Kusari et al. 2012; Panaccione et al. 2014), and mediating plant functional trait expression (Griffin et al. 2016; Friesen et al. 2011). Recent work has demonstrated how these various effects of endophytes can influence whole ecosystem level processes (Christian et al. 2019; Clay and Holah 1999; Griffin et al. 2017; Kivlin et al. 2013; Laforest-Lapointe et al. 2017b). Importantly, endophytes are often erroneously assumed to have predominantly mutualistic associations with their hosts. Reality is much more complex and the influence of endophyte taxa is highly context dependent (Carroll 1988; Rodriguez et al. 2009), with interactions between hosts and endophytes ranging from mutualism through commensalism to latent or mild antagonism (Hardoim et al. 2008; Schulz and Boyle 2005; Saikkonen et al. 1998).

Much of the focus on endophytes has been driven by applied scientists interested in harnessing these taxa as a means to manipulate plant phenotype (e.g., increase growth; Doty 2011) and prevent pathogen colonization of crops (Busby et al. 2017). Endophytes have also attracted attention from natural products chemists who survey the world’s organisms for useful compounds (Aly et al. 2010; Strobel and Daisy 2003). This is motivated by the capacity of various endophytes to synthesize an impressive array of bio-active small molecules (Newman et al. 2003; Strobel et al. 2004; Verma et al. 2009). Indeed, a number of endophyte-synthesized compounds are of medicinal value (Kharwar et al. 2011; Strobel et al. 1996).

Both basic and applied research regarding endophytes have been hampered by a lack of knowledge regarding the distribution of endophyte biodiversity at any spatial scale—from global, inter-biome scales to among the tissues of individual host plants. Similarly, almost nothing is known regarding how endophyte biodiversity maps onto the plant phylogeny. Characterizing such broad patterns in endophyte biodiversity is extremely logistically challenging because of the outlay of effort required for large culturing and sequencing projects (Arnold and Engelbrecht 2007; Carini 2019; Nilsson et al. 2018). Moreover, determining the causes of patterns in endophyte diversity is difficult because they result from the interplay of many forcings, including both contemporary and historical ecological drivers (i.e., niche determination, ecological drift, dispersal limitation) and, at longer timescales, evolutionary processes (i.e., divergence and extinction; Hanson et al. 2012; Wiens and Donoghue 2004).

Further complicating matters is the disconnection between the spatial scale of sampling endophyte assemblages, as dictated by logistical constraints, and the size of the focal organisms. This problem was well illustrated by Remus-Emsermann and Schlechter (2018) who point out that the disparity in size between a single bacterium of 2 µm^3^ and a leaf mirrors the ratio of sizes between a person and a mid-sized country. Indeed, using traditional culturing and sequencing methodologies, we can only sample what are in effect whole regions of endophytes that may include multiple assemblages that never directly interact, that have been shaped by differing community assembly processes, and that may even have divergent evolutionary histories. This complicates the study of endophyte biogeography and community ecology because the scale of sampling is so much larger than many covariates that may affect membership of endophytes in a particular assemblage. For instance, microhabitat variation within leaves (such as proximity to upper or lower leaf surfaces, veins, etc.) may have effects on endophyte assemblages akin to those of shifting elevation across a mountainside on forest composition, and those forcings are unavailable for study when the unit of replication is an entire leaf, or even a leaf section (Herre et al. 2007; Lodge et al. 1996; Remus-Emsermann and Schlechter 2018; Vacher et al. 2016). Adding further complexity, some bacterial endophytes can live inside endophytic fungi (Shaffer et al. 2016); thus, for these bacteria, the habitat covariates most relevant for explaining inter-assemblage variation may be the traits of the host fungus, not the traits of the host plant.

While we have much to learn regarding the distribution of endophyte biodiversity, patterns observed so far generally follow the predictions of community ecology theory and are similar to those observed for metazoan and plant assemblages (Christian et al. 2015; Nemergut et al. 2013). For instance, sampling of endophyte assemblages recapitulates the positive species-area relationship observed in so many natural systems—as one samples a larger area one encounters more taxa (e.g., Suryanarayanan et al. 2002). A necessary correlate of this observation is that the similarity among endophyte assemblages declines with distance, which also has been demonstrated numerous times (e.g., Davis and Shaw 2008; Higgins et al. 2014; Nemergut et al. 2013; Vacher et al. 2016). While its causes are multifarious and poorly understood (Martiny et al. 2006; Vellend 2010), the existence of distance-decay suggests that endophytic microbes are influenced not only by deterministic forcings, but also by what are typically regarded as neutral processes, such as dispersal limitation and ecological drift (Hubbell 2001; MacArthur and Wilson 2001; Nemergut et al. 2013). Also following what is known for most large, multi-cellular organisms, foliar fungal endophyte biodiversity seems to follow a latitudinal gradient, with higher diversity at lower latitudes, as shown by Arnold and Lutzoni (2007). The reasons for this pattern remain unknown, but likely include both contemporary and historical drivers (Mittelbach et al. 2007; Pianka 1966). Importantly, it is unclear if this pattern holds for non-fungal taxa and if non-foliar tissues harbor higher fungal endophyte richness at lower latitudes. Indeed, ectomycorrhizal fungi appear to be at their richest in temperate zones (Tedersoo et al. 2014), which suggests the possibility that other root associated microbes may be richest at intermediate latitudes as well.

Much of what is known regarding patterns of endophyte biodiversity demonstrates the influence of contemporary ecological contingencies at either regional or local spatial scales (i.e., niche determinism). For instance, Zimmerman and Vitousek (2012) reported greater fungal endophyte richness at wetter, low elevation sites on a Hawaiian mountainside and Bowman and Arnold (2018) found that *Pinus ponderosa* hosted more diverse foliar fungal endophyte communities at mid-to-high elevations compared to lower elevations in southwestern Arizona (also see Giauque and Hawkes 2013; Glynou et al. 2016; Lau et al. 2013). Furthermore, it is clear that endophyte assemblages shift among coexisting host species, though the effect of host on endophyte assemblage divergence can be quite modest in some cases (Griffin et al. 2019; Redford et al. 2010; Vincent et al. 2015). While evidence reported to date suggests that many cultivable endophytes are host generalists (e.g., Arnold and Lutzoni 2007; Suryanarayanan 2018), specialist endophytes do exist, as demonstrated by the fidelity of vertically-transmitted (seed borne) *Epichloë* fungi to members of Poaceae (Clay and Schardl 2002; Rudgers et al. 2009) and of swainsonine-producing *Alternaria* fungi to certain Fabaceous taxa (Cook et al. 2014; Panaccione et al. 2014). However, the host range of the myriad endophytes that occur at low relative abundances is unknown (Arnold and Lutzoni 2007)—an important gap in knowledge given that these taxa likely comprise the bulk of endophyte biodiversity (Arnold and Lutzoni 2007; Lynch and Neufeld 2015).

Even within an individual plant, niche determinism can shape endophyte assemblages as many studies have confirmed that endophyte assemblages vary among tissue types (e.g., the endophyte assemblages in roots often differ from those in leaves; e.g., Coleman-Derr et al. 2016; but see Haruna et al. 2018; Massoni et al. 2019), though general patterns in endophyte richness among tissue types have not been described.

Niche determinism is not the only force affecting endophyte biodiversity within a particular substrate, though it is likely the best studied. For instance, it is clear from research within non-endophytic taxa that microbes can be dispersal limited and therefore the long-standing Baas Becking hypothesis for microbial biogeography, namely that “everything is everywhere, but the environment selects” (Baas Becking 1934) is too simplistic (Hanson et al. 2012; Martiny et al. 2006; Nemergut et al. 2013; Vellend 2010). It is still unclear, however, how dispersal limitation and ecological drift (stochastic changes in community membership due to aggregated life history events) shape endophyte community assembly. Similarly, little is understood regarding the influence of historical factors, including divergence and extinction, on endophyte biogeography, though it seems likely that these factors will manifest in differences in endophyte assemblages across biomes and geographical regions, just as they do for other taxa (Mittelbach et al. 2007). How often these forces act at what are, to our sensibilities, small spatial scales, such as within a forest or even a single-long-lived tree, remains unknown.

A final challenge to the study of endophyte distribution is the likelihood that most patterns in biodiversity will be taxon specific, because taxa respond differently to ecological contingencies and are on varying evolutionary trajectories. For example, Coleman-Derr et al. (2016) suggest that prokaryote taxa are more influenced by plant tissue type than fungal taxa, which are affected more by host habitat and biogeography. At the order level, Jumpponen et al. (2017) suggested that Helotiales fungal root endophytes are most abundant in forested ecosystems and Pleosporales fungi are more common in grasslands. Studies such as these are very rare; very little is known regarding the geographical or host ranges of endophytes at any level of the biological hierarchy—from phylum to subspecies.

While daunting, the study of endophyte biogeography and community assembly will likely provide important benefits for both basic and applied research. An exemplar is provided by Higginbotham et al. (2013) who isolated over 3000 endophytic fungi from numerous tropical angiosperms and ferns and tested these cultures against common diseases, including malaria, Chagas disease, and cancer. They report that 30% of the fungi showed strong activity against at least one of the focal diseases and that bioactivity against a specific target was non-randomly distributed across the fungal phylogeny. Intriguingly, they also reported a generally higher degree of bioactivity in taxa sourced from cloud forests compared to lowland tropical forests—thus providing a biogeographic road-map for natural product discovery in tropical forests that demonstrates an important role of both biome and host phylogeny (also see Schulz et al. 2002).

The first step towards a working knowledge of endophyte distributions across spatial scales is the description of broad patterns in their biodiversity. To understand the scope of relevant research, we scoured the literature and extracted basic metadata from 596 studies characterizing endophyte assemblages. Our goal was to synthesize the meta-data from representative studies, with the hopes of highlighting particular portions of the plant phylogeny and specific biomes that need further exploration and to determine how information could be shared among studies. Additionally, we paired our survey with a vote counting procedure where we compared patterns of endophyte richness among tissue types. The synthesis process illustrated the challenges of pooling information among studies and, consequently, we offer specific guidelines for data sharing and research reproducibility moving forward. For our purposes in this article, we did not consider obligate pathogens, epiphytes, or mycorrhizae; nor did we include a review of the large body of literature examining *Rhizobia* and their associations with legumes, as others have already done so (e.g., Peter et al. 1996; Willems 2006).

## Methods

We searched Google Scholar and Web of Science for the term “endophyte” in conjunction with “fungal”, “bacterial”, “diversity”, or “community”. All publications in which the authors characterized endophyte assemblage biodiversity were collated. As we were primarily interested in studies characterizing endophyte biodiversity, we did not consider research involving manipulative experiments where no survey of microbial diversity was conducted. We also made the choice to omit studies that did not distinguish between epiphytes and endophytes through performing some form of surface sterilization. Searches were performed periodically from 2016–2018 and additional studies added to our database as we became aware of them until the beginning of 2019. We apologize to those authors whose work we missed and to those who have published their work in non-English language journals, which typically did not appear in our searches and were inaccessible to us because of our linguistic backgrounds.

From each study, we collected information on host organism(s) studied, research location(s), tissue type(s) surveyed, and various metadata describing the nature of the survey conducted—for instance, if the endophyte assemblage was characterized via sequencing or culturing, if spatial or temporal replication was employed, host and culture vouchers deposited, and data made available. We considered studies spatially replicated if they involved two or more sampling locations separated by *≥*1km. We chose this threshold because of work by Higgins et al. (2014) who reported rapid distance-decay in endophyte assemblage similarity within tropical grasses within 1 km. It is likely that the strength of distance-decay depends upon biome, host plant, endophyte taxon, and other ecological conditions, thus determining what constitutes sufficient spatial replication is challenging and study-dependent. We use a 1 km threshold here because most studies that were replicated at a smaller spatial scale were within a single field, forest, or greenhouse and thus were likely exposed to similar endophyte inoculum, at least over longer timescales. If the study location was not explicitly provided, we extrapolated an estimate based on the city or country reported by the authors. We assigned studies to biomes following the nomenclature of Olson et al. (2001). Host plants collected from urban, agricultural, or areas that were otherwise managed, were classified as coming from “cultivated” landscapes and counted independently from those studies that occurred in unmanaged landscapes within the same biome. We chose this approach because managed areas experience ecological contingencies divergent from their surroundings (e.g., irrigation). We considered studies of “stems” as those involving sampling of woody branches, twigs, or grass shoots. Studies of “roots” included any survey of below-ground plant tissue, but excluded rhizosphere soil surveys and studies which did not attempt to surface sterilize roots. We considered studies of “leaves” to be those sampling leaf sections or whole leaves/leaflets (including needles) and did not consider studies that combined petioles with leaf blade tissue.

To understand the phylogenetic breadth of host plants surveyed, we calculated the total number of hosts examined for each plant family and plotted this information on a phylogeny of the Embryophyta (algal endophyte hosts were thus omitted from this portion of our analysis) generated using phyloT (online software accessible at https://phylot.biobyte.de/). The National Center for Biotechnology Information taxonomy database was used to generate the tree (database accessed March 15, 2019; Federhen 2012). iTOL v4.3.2 was used for tree visualization (Letunic and Bork 2016). All data manipulation was performed in the R statistical computing environment (R Core Team 2019).

### Vote counting to determine patterns in relative tissue richness

In addition, we asked how endophyte richness shifted among tissue types, for both fungi and bacteria. We attempted two approaches to address this question—a formal meta-analysis and a simple vote counting approach. Because few studies used the same methods, comparing the effect of tissue type on richness *among* studies was inappropriate (this limitation also precluded comparison of richness among taxa or across biomes, unfortunately). Thus, we only examined those studies that compared richness among multiple tissue types and all comparisons were made *within* studies.

Unfortunately, very few studies provided data sufficient for a quantitatively rigorous meta-analysis (see Supplemental Methods and Results), so we conducted a simple vote counting procedure where we considered each study independently and ranked tissue types by the relative richness reported in that study. We only considered those studies that examined multiple tissues and that standardized sampling effort among tissues. In total, we examined 243 studies: 182 studies of fungal endophytes and 61 studies of bacterial endophytes. After ranking tissues by relative richness separately for each study, we calculated, across studies, the proportion of times one tissue type had higher richness than another tissue (e.g., for what proportion of studies did leaves have higher richness than roots) and calculated the probabilities of these proportions using a binomial sign test (Cooper and Hedges 1993). This test is simply the probability of observing a particular number, or more, of positive outcomes (in our case, one tissue type having higher richness than another) given a certain number of trials and assuming equal probability of positive and negative outcomes. For this vote counting approach, we focused on richness because fewer studies reported diversity metrics and, when not explicitly reported by authors, relative richness was simpler to calculate and extract from published summary tables and figures than were diversity entropies. To test how growth habit influenced relative microbial richness among tissues, we conducted vote counting separately for studies of hosts with the following growth habits: woody-stemmed trees and shrubs, forbs, and graminoids.

## Results

Our survey highlighted the breadth of the endophyte biodiversity literature, as we extracted data from 596 unique publications. We report that interest in endophyte diversity is on the rise, with a sharp increase in studies per year since 2010 (Fig. 1). Fungi have received comparatively more attention than bacteria, though this disparity is diminishing (Figs. 1 &2e). The majority of studies were of foliar endophytes (1694 unique combinations of study and host species), followed by root (577 combinations) and stem (540 combinations) endophytes. By comparison, floral tissues (39 combinations) and plant propagules were understudied (172 combinations; Fig. 2). Multiple-host studies were not the norm—approximately ∼66% of studies focused on a single host taxon.

**Figure 2:**
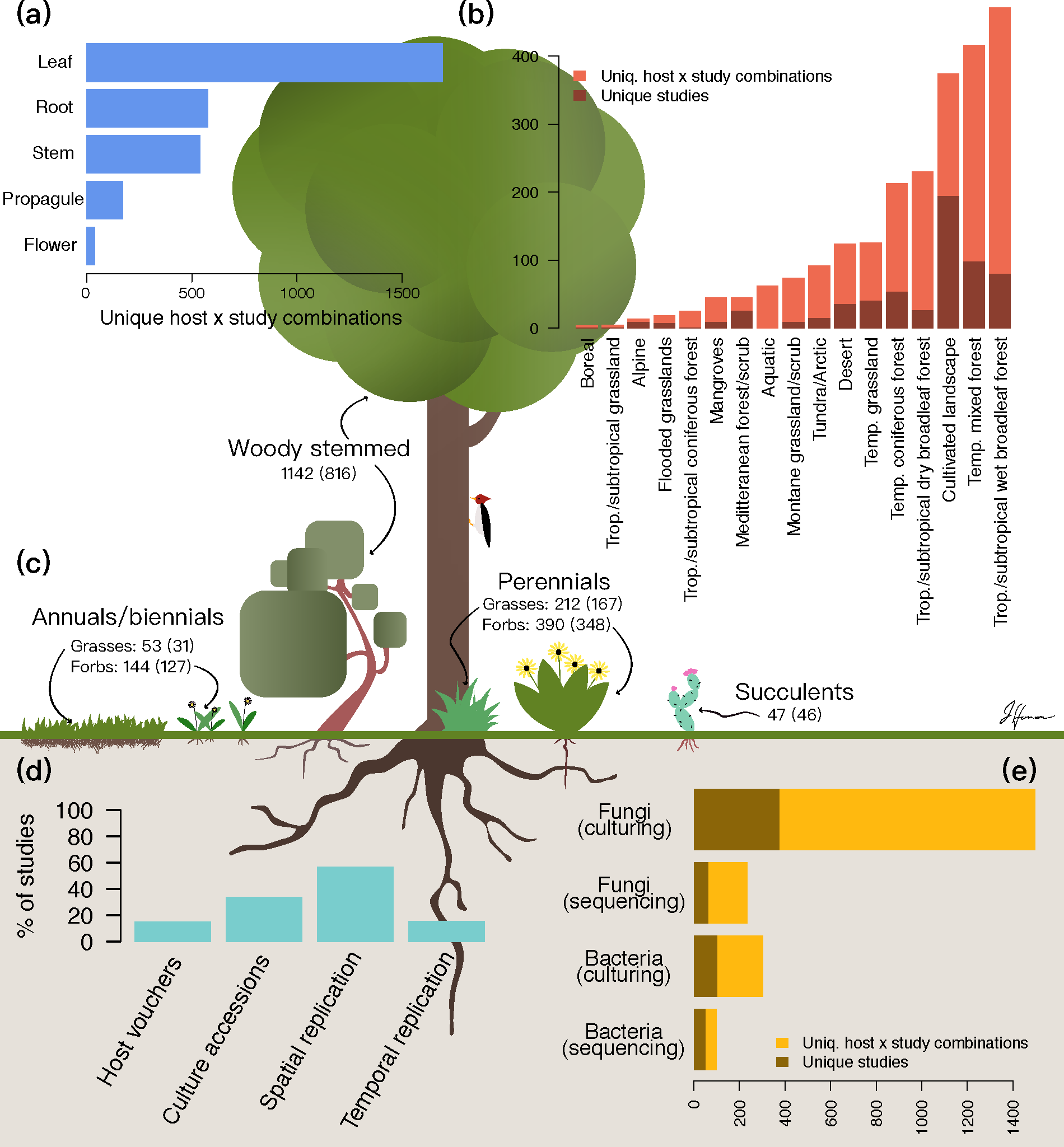
Summary of 596 publications characterizing endophyte biodiversity. Because many studies surveyed multiple hosts, we report both number of studies and number of unique host by study combinations. We counted the number of studies surveying each plant compartment (a), biome (b), and host life history category (c; values in parentheses are unique hosts). We also extracted information pertaining to study design and reproducibility (d). Finally, we determined the endophytic taxon characterized and the methodology employed (e).

### The global extent of endophyte biodiversity research

The geographical range encompassed by the studies we considered was global; endophytes, both fungal and bacterial, have been recovered from hosts across all major biomes (Fig. 3). Temperate mixed coniferous and deciduous forests were the best studied, with 98 studies (16% of total). However, the most unique combinations of host and study were reported from tropical and subtropical wet forests (471, 21% of total). This was due to several studies that surveyed many hosts within these forests (e.g., Rojas-Jimenez et al. 2016 with 92 hosts and Suryanarayanan et al. 2011 with 70 hosts). In terms of unique studies, research in tropical and subtropical forests composed a more modest 13% of studies in our survey. Many biomes were quite understudied. For instance, 50 or fewer studies (in terms of unique host by study combinations) were conducted in seven of the seventeen biomes that we considered (Fig. 2b). Together, studies from these biomes composed only 7% of those surveyed.

**Figure 3:**
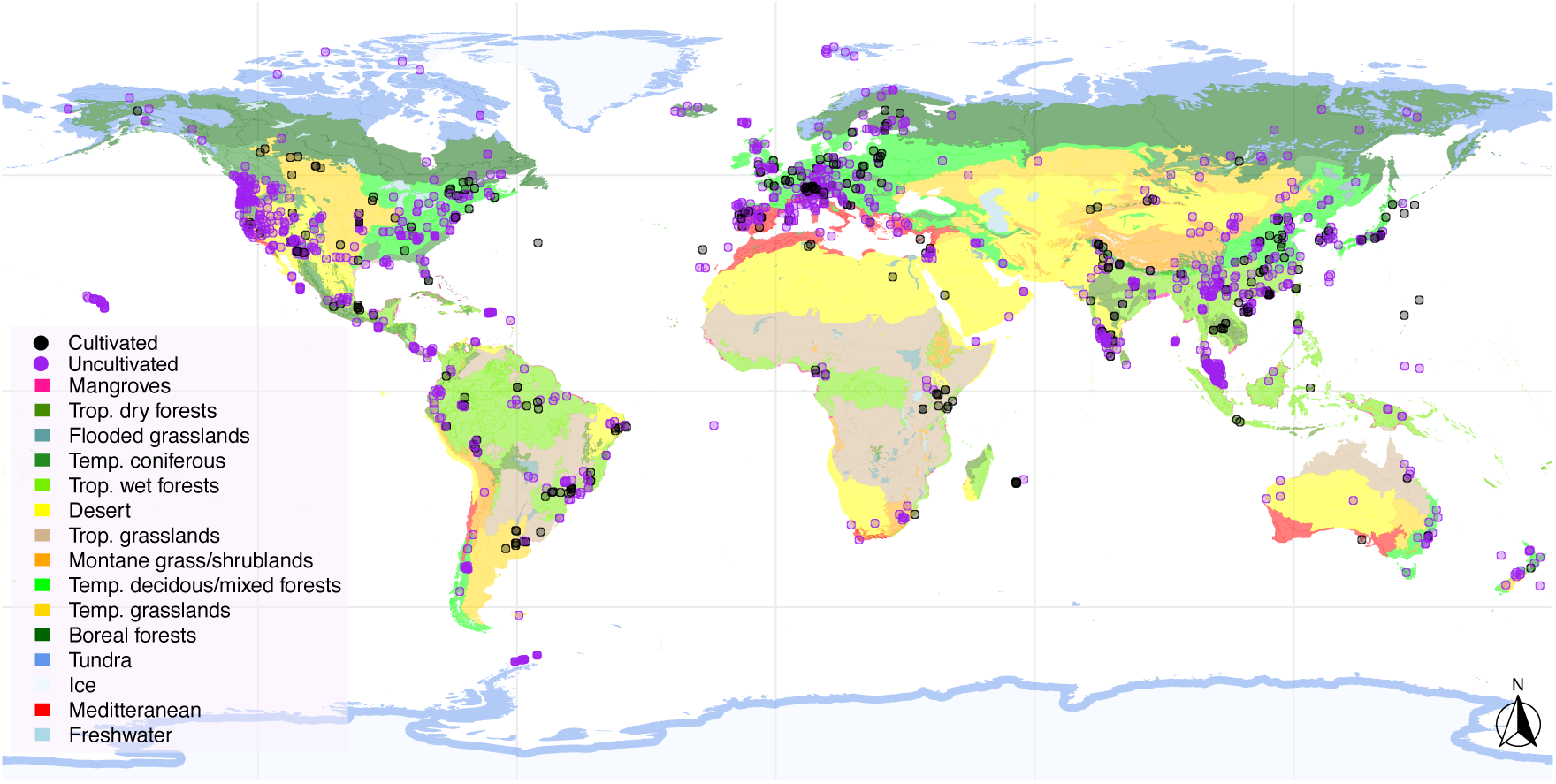
Locations of studies considered. An interactive, zoomable version of this map can be found at: https://jharrisonecoevo.github.io/EndophyteMap/. Black points represent studies of cultivated crops and managed landscapes, purple points represent studies of uncultivated plants in unmanaged settings. Biomes are color coded and delineated in accordance with Olson et al. (2001). In some cases, multiple, proximal locations were surveyed and a single point was used to graphically represent these locations. If a study did not provide exact location information then study location was approximated.

Across biomes, we found comparatively few studies of hosts growing in obvious wilderness, far from human development. Indeed, 33% of studies relied on hosts grown in cultivated environments, including urban locations, agricultural landscapes, and greenhouses (with university campuses being particularly well sampled). This estimate may be conservative as for some studies the exact collection location was difficult to determine and so we did not include them in the “cultivated” category, but sampling was likely not far from human development.

### Much of the host phylogeny remains unsampled

The studies we surveyed encompassed 1702 unique taxa from 254 plant families. Poaceae was by far the most well-studied family (189 hosts studied), followed by Fabaceae (98 hosts), Pinaceae (82 hosts), and Asteraceae (79 hosts; Fig. 4). In the studies we examined, fungal endophytes have been surveyed in hosts from 30% of plant families listed in the NCBI taxonomy database for Embryophyta. By comparison, bacterial endophytes have been characterized in only 10.5% of plant families. Of particular note, very few observations of foliar microbiota have been made among bryophyte and pteridophyte families (Fig. 4; Davis and Shaw 2008; Desirò Alessandro et al. 2013). Additionally, we observed a mismatch between host family species richness and sampling effort. For instance, only 29 Orchidaceae species have been surveyed out of the approximately 28,000 accepted orchid taxa occurring worldwide (The Plant List, Chase et al. 2015).

**Figure 4:**
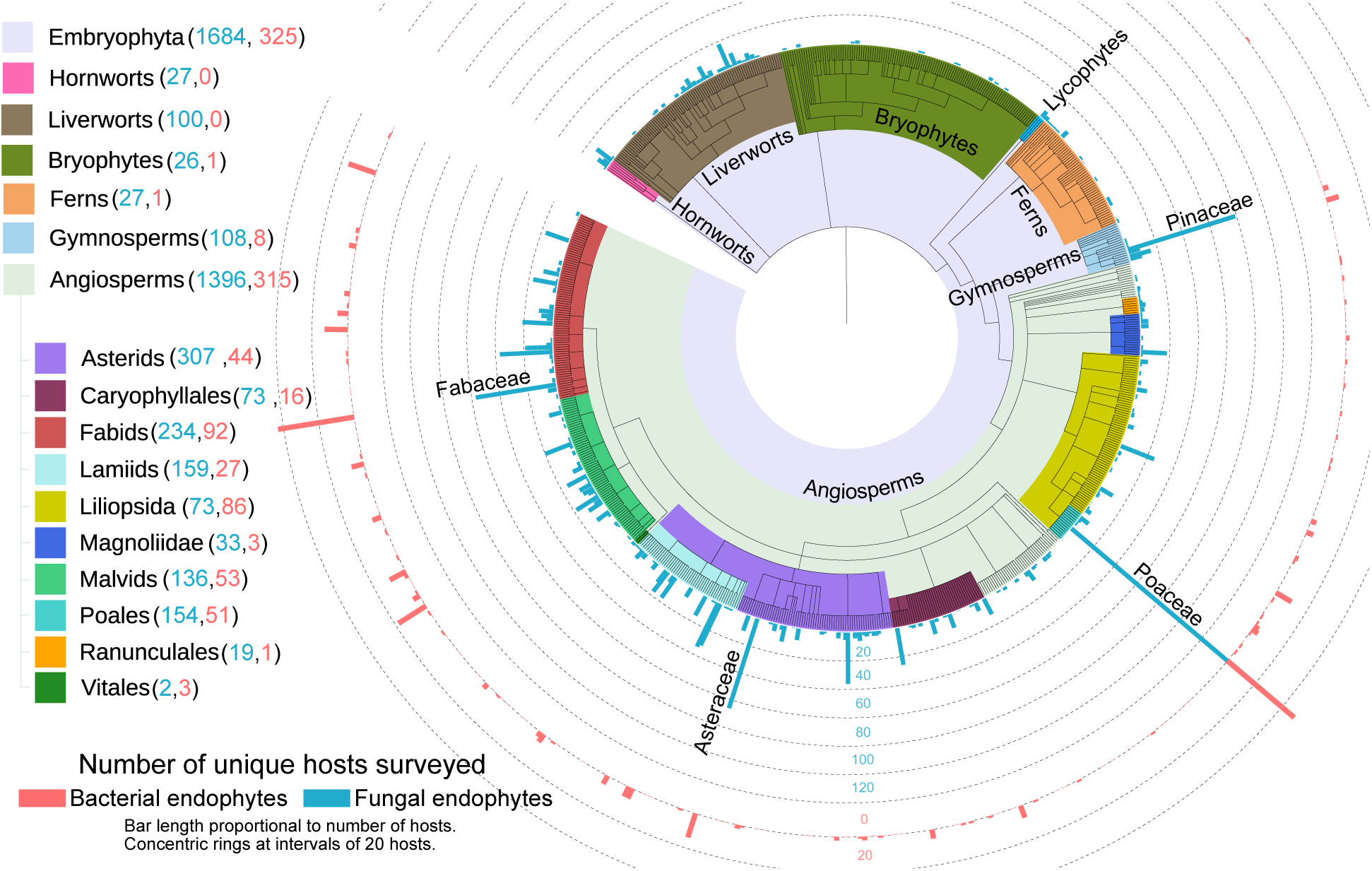
Survey effort across Embryophyta. Number of studies surveying fungal (blue) and bacterial (red) endophytes are shown extending outwards from the tips of the phylogeny. Tips are families. Notable taxa within Embryophyta are labeled and color-coded. Numbers in parentheses denote unique hosts surveyed. Very few surveys of bacterial endophytes have been conducted in bryophyte hosts, therefore this portion of the figure has been abbreviated to aid visualization. An interactive, zoomable version of this phylogeny can be found at: https://itol.embl.de/shared/harrisonjg.

### Replication and reproducibility could be improved

We also characterized details for each study regarding sampling scheme and reproducibility (Fig. 2de). We found that just over half of studies were spatially replicated (sampling areas were separated by at least a km) and fewer than a quarter of studies were temporally replicated. The majority of studies (∼74%) relied on culturing, however only about a third of these studies reported accessioning cultures (Fig. 2e). By comparison, 72% of studies that relied on sequence data provided clear instructions for downloading raw data, though only 23% of these studies provided processed data (such as an OTU table). Surprisingly, fewer than 20% of studies mentioned accessioning host vouchers. For cultivated plants, we considered a description of the cultivar as equivalent to an accessioned voucher.

### The effects of tissue type on endophyte richness and diversity

We performed vote counting to compare the relative richness and diversity of fungal endophyte assemblages in varying tissue types across plant taxa. We resorted to vote-counting because data were insufficient for a robust meta-analysis (see Supplemental Methods and Results). We found that relative tissue richness was dependent upon host growth habit. For instance, stems had richer fungal endophyte assemblages than leaves for woody-stemmed hosts, but this pattern was not observed for either forbs or graminoids (Table S1). By comparison, for graminoids, roots had richer fungal and bacterial endophyte assemblages than stems (Table S3). For forbs, no tissue type was clearly richer, on average, than other tissues (Table S2). Additionally, for fungal endophytes, we found that reproductive structures, including flowers and propagules, were relatively species poor, while bark was species rich (Table S1 &S2), though these results are quite tentative given the few studies that compared endophyte assemblages in these tissues to those in other portions of the plant corpus.

## Discussion

We report that endophyte biodiversity has been studied within all major biomes and continents (even Antarctica, if one counts King George Island; Fig. 3; Rosa et al. 2009). Given that widespread interest in endophytes did not occur until the 1970s, progress has been rapid. However, great swathes of the globe still remain unsurveyed. Certain biomes have been particularly understudied—either due to their high biodiversity, which makes thorough sampling exceptionally difficult (i.e., tropical rainforests); large geographical area (e.g., the boreal forest); or because they are geographically restricted and simply have not received much attention. For instance, we found few studies from coastal dunes, flooded grasslands, and mangrove forests. These habitats are challenging for plants, due to salinity, short intervals between disturbances, and the presence of anoxic soil. Surveys of understudied biomes will help define the scope of endophyte biodiversity and functional traits. In particular, we suggest that surveys in flooded grasslands and mangroves may improve our understanding of archael endophyte biodiversity (Moissl-Eichinger et al. 2018), as this branch of life includes numerous halophiles and other extremophiles that may be able to cope with the abiotic conditions characteristic of those locations. Similarly, studies in desert and alpine biomes may uncover endophytes with unique mechanisms for coping with the severe ultraviolet exposure, temperature swings, and desiccation that occurs in such harsh habitats (see for example, Lopez et al. 2011; Massimo et al. 2015; Sangamesh et al. 2017).

We also reported a lack of studies from Africa, west and north Asia, and the interiors of Australia and South America (Fig. 3). These areas hold some of the most biodiverse and charismatic landscapes on the planet; for instance, the Congo basin is the second largest tropical rainforest in the world, with thousands of endemic plant taxa (Brenan 1978; Linder 2001), and it has historically experienced less deforestation than other rainforests (Koenig 2008). Similarly, the Cape Floristic province in Africa has some of the highest levels of plant endemism in the world. Because these regions have evolutionary histories that have facilitated endemism, it seems likely that they harbor unique endophyte taxa and would be prime locations to study coevolution and codivergence between plants and endophytes. More generally, the lack of sampling outside of North American, Europe, and portions of Asia precludes a robust knowledge of endophyte biogeography.

### The influence of human development on endophyte biodiversity

We acknowledge the logistical challenges of sampling the remote locations that remain understudied. Indeed, we report an imprint of this challenge in even relatively well-studied regions, where we found that most studies were conducted near roadways, townships, and other human development. The lack of sampling in wilderness areas likely biases our nascent understanding of endophyte biodiversity. Human development is associated with pollution, habitat fragmentation, ecosystem disturbance frequency, and the abundance of introduced hosts (Crowl et al. 2008; Dietz et al. 2007)—all of which likely affect plant microbiomes. Evidence for this hypothesis is sparse, however Laforest-Lapointe et al. (2017a) reported many phyllosphere bacterial taxa shift in relative abundance along an urbanization gradient, with an overall decline in dominant Alphaproteobacteria with more urbanization. Similarly, Lappalainen et al. (1999) reported a decline in endophyte colonization of *Betula* trees with proximity to a copper-nickel smelter. Variation in heavy metal concentrations (Tóth et al. 2009; Jurc et al. 1996), acid rain (Helander et al. 1994), and air pollution (Wolfe et al. 2018), have all been associated with shifts in endophyte assemblages—thus, it seems likely that the effects of pollution and urbanization are multifarious and have effects which depend upon the endophytic taxon examined and the ecological context.

In addition to pollution, habitat fragmentation also increases in proximity to human development. Very little is known regarding how habitat fragmentation affects microbial assemblages or, more generally, how metacommunity processes manifest within microbiomes (Christian et al. 2015). However, classic island biogeography theory (MacArthur and Wilson 2001) suggests that human-caused habitat fragmentation likely shapes endophyte assemblages through determining proximity to inoculum sources. In a survey spanning islands of various sizes, Helander et al. (2007) reported that endophyte colonization of *Betula* spp. trees was greater on larger islands and islands closer to the mainland (also see Oono et al. 2017). This result, coupled with work documenting dispersal limitation in non-endophyte, microbial systems (Golan and Pringle 2017; Peay et al. 2010; Peay et al. 2007; Andrews et al. 1987) suggests that it is reasonable to expect variation in endophyte assemblages routinely follows the predictions of island biogeography, regardless of whether habitat fragmentation and patch size is caused by geological processes or human influence.

Another way in which endophyte assemblages may be affected by proximity to human development is through the influence of invasive plant taxa, which are often much more abundant near development than in wilderness areas. Invasive host taxa could influence endophytes in a variety of ways—from changing the inoculum pool within an area (i.e. “neighborhood” effects; Moeller et al. 2015), bringing along endophyte taxa or genotypes from the ancestral range of the host (Dickie et al. 2017), or affecting many other aspects of the local ecology (e.g. shifting fire regimes [Brooks et al. 2004], determining litter deposition rate and elemental composition [Allison and Vitousek 2004], influencing herbivore assemblages [Forister 2009], etc.).

Interestingly, the few studies we encountered that surveyed endophytic microbiomes of invasive plants in both their native and invaded ranges found that endophyte assemblages differed between ranges. For instance, Lu-Irving et al. (2019) report reduced richness in phyllosphere and endophytic root bacteria in the invaded portion of the range of *Centaurea solstitialis* compared to the native range. Similarly, Shipunov et al. (2008) report a wholesale shift in the fungal endophyte assemblage within the leaves of *Centaurea stoebe* in invaded versus native portions of its range. Thus, the biodiversity of endophytes within widespread, invasive plants is also influenced by host invasion history (also see Gundale et al. 2016; Sikes et al. 2016; Taylor et al. 1999).

All of these anecdotes support the idea that endophyte assemblages in relatively undisturbed areas, such as portions of the Amazon or the Siberian forest, are likely to be different from those in conspecific hosts growing near human habitation or that are being actively cultivated and thus remote locations should be the focus of further study. Even if different microbial taxa are not observed in less disturbed environments, study of the shifts in relative abundances among endophyte assemblages along urbanization and pollution gradients could provide insight into how endophytes interact and communities assemble (e.g., Gazis and Chaverri 2015).

### Much of the host phylogeny remains unexplored—what might we be missing?

We found that members of about a third of plant families have been surveyed for fungal endophytes and only about a tenth of plant families are represented among studies characterizing bacterial endophyte assemblages. These results suggest we may be missing a large portion of endophyte biodiversity. It is true that many cultivable endophytic taxa are known to have broad host ranges (e.g., Arnold and Lutzoni 2007; Suryanarayanan 2018), thus one could argue that an understanding of endophyte biodiversity does not hinge on thorough sampling of potential host taxa. However, we note that, in the majority of multivariate studies of endophyte biogeography, host taxon is an important predictor of assemblage variation (Griffin et al. 2019; Kivlin et al. 2019)—albeit a sometimes modest one (Vincent et al. 2015). Moreover, little is known regarding the host range of those rare endophyte taxa that compose the bulk of most assemblages (Arnold and Lutzoni 2007).

Studies delineating host range are desperately needed to understand endophyte distributions and biodiversity, however given the daunting nature of the sampling required, where then should we begin? We suggest targeting those plant lineages with unique traits, such as production of unusual secondary metabolites or preferences for restricted or harsh habitats (e.g. halophiles and extremophiles). As an example, certain *Astragalus* taxa can hyperaccumulate selenium, and recent research has suggested that these plants may harbor unusual endophytic taxa that could influence selenium uptake (Sura-de Jong et al. 2015; Lindblom et al. 2018; Lindblom et al. 2013). Following a similar rationale, we also suggest surveying those plant families that are phylogenetically distinctive. If coevolution or codivergence has occurred between hosts and their endophytes, then unusual endophytic taxa could occur in hosts from remote portions of the plant phylogeny (Hassani et al. 2019). Non-vascular plants, in particular, deserve more attention, as these plants have different evolutionary histories, physiology, growth habits, and preferred habitats than vascular plants (Huang et al. 2018).

An additional justification for surveying broadly across the plant phylogeny is the discovery of specialist endophyte taxa. Surveys of seeds, in particular, could lead to the discovery of more vertically-transmitted endophytes (class I and II endophytes *sensu* Rodriguez et al. 2009), which are particularly interesting because of their capacity to influence their hosts during early ontogeny (e.g., Gundel et al. 2017; Hodgson et al. 2014; Truyens et al. 2015). An individual seed generally contains a very species poor endophyte assemblage (e.g., in many cases only a single fungus was isolated from seeds, see Hodgson et al. 2014; Newcombe et al. 2018; Shipunov et al. 2008), and relatively few instances of vertical transmission of endophyte taxa have been documented. However, recent work by Hodgson et al. (2014) provides evidence that vertical-transmission of fungi may occur much more often than previously suspected (also see a review on bacterial seed endophytes by Truyens et al. 2015). Indeed, while the well-known clavicipitaceous endophytes seem to be limited to members of the Poaceae (Rudgers et al. 2009), the occurrence of vertically-transmitted endophytes capable of systemic growth has been documented from throughout the plant phylogeny, including within members of the Asteraceae (Hodgson et al. 2014), Araliaceae (Soares et al. 2016), Convulvulacea (Cook et al. 2013), Ericaceae (Rayner 1915), Fabaceae (including members of *Astragalus, Oxytropis* and *Swainsona* Cook et al. 2009; Cook et al. 2014; Grum et al. 2013), Papaveraceae (Hodgson et al. 2014), and Plantaginaceae (Hodgson et al. 2014). This suggests that facultative vertical-transmission may occur in numerous plant hosts and across many biomes. Cross-biome comparative studies of the seed microbiome could determine if vertical-transmission is more common in certain habitats, as might be predicted if these endophytes interact mutualistically with their hosts to ameliorate the negative affects of particular abiotic conditions (Afkhami et al. 2014; Gundel et al. 2017).

### The effects of tissue type on endophyte assemblages

Our vote counting approach suggested that in woody plants stems had higher richness than other tissues, for both fungi and bacteria. However, for graminoids, roots were the richest tissue and, for forbs, inter-tissue patterns in richness were less clear (Tables S1–S3). These results suggest that tissues with greater lifetime inocula exposure have the highest richness across plant life histories. Indeed, several studies have demonstrated that older leaves typically harbor richer microbial assemblages than younger leaves, presumably because of greater exposure to inoculum and increased time for microbial growth (Arnold et al. 2003; Ercolani 1991). Stems and bark of woody plants are exposed to inocula in air, water, and dust year round and have long lifespans (indeed much bark is dead and can remain on the trunk for a lengthy period of time), whereas leaves, even for evergreen trees, do not persist for nearly as long. Similarly, roots are the longest-lived tissues of many perennial forbs and graminoids, as above-ground tissues of these hosts often senesce annually. It is true that roots of woody-stemmed plants can be quite long-lived, however roots are primarily encountering inoculum from the surrounding soil matrix, thus it is possible that there is greater variation in the inoculum encountered by stems than by roots over the lives of those tissues. Alternatively, perhaps the resources available to microbes within stems of woody-plants favor higher richness compared to leaves, particularly of latent saprotrophs that catabolize lignin or other structural carbohydrates (Oses et al. 2006; Oses et al. 2008). These hypotheses are not mutually exclusive and await experimental testing.

Our survey comes with several caveats. First, it is possible that the efficacy of surface sterilization may vary with tissue type; thus, for instance, the high fungal richness in bark that we report could be because it was more difficult to surface sterilize than other tissues. Also, while we chose those studies that had the same sample size (in terms of replicates) between each tissue type, it was not always apparent that the same mass was used for each sample. Additionally, both culture and sequence-based surveys suffer from taxonomic biases (Carini 2019; Nilsson et al. 2018) and if those biases coincide with taxonomic variation among tissue types, then richness estimates will be incorrect. Nevertheless, our analysis demonstrates the existence of clear patterns in richness among tissue types and suggests several hypotheses for those patterns that deserve further study.

### How can we best share information among studies?

We report several challenges that impede meta-analysis and synthesis of the endophyte literature (e.g., Meiser et al. 2014). Most importantly, raw and processed sequence data were not always available. Moreover, it was quite rare for sufficient detail to be provided regarding sequence processing—including options and versions for software used and date accessed for taxonomy training databases, which are in constant flux. Given the challenge in reprocessing data and the influence different bioinformatic pipelines can have on results (e.g., Pauvert et al. 2019), we suggest that publication of polished data and scripts should be considered to facilitate information sharing among studies. Those data that would be most amenable to meta-analysis include replicate by taxon tables, sequences of operational taxonomic units (OTUs) or exact sequence variants (ESVs), and the taxonomic hypotheses for those sequences. In many cases, meta-analysis will require substantial reprocessing of the data, so raw data should also be made available.

Additionally, we suggest that authors consider depositing vouchers of host taxa studied, nucleic acids extracted, or cultures obtained, in an herbarium whenever possible (Fig. 2d). This suggestion is motivated, in part, by fascinating new work by Daru et al. (2019) who have shown that endophytes within herbarium specimens can be sequenced, and, in some cases, even cultured. Thus, vouchers could act as “time capsules” that preserve endophyte genotypes and could afford insight into endophyte evolution and shifts in host and geographic range over time. To best share information among vouchers, standardized protocols (such as drying time and temperature) could be helpful to adopt, though we acknowledge the challenge of implementing such standards during field collection. Deposited cultures could provide many of the same benefits as host vouchers, but would also allow researchers to grow endophytes of interest to meet various experimental goals (Huang et al. 2018; Suryanarayanan 2019). Finally, the plant taxonomy is ever-changing, thus as future researchers interpret published work, they may wish to examine accessions to determine the most current taxonomic placement of the focal host or endophyte. In sum, we see herbaria as tremendous resources for the study of the plant microbiome, and, consequently, we urge participation in their continued development.

## Conclusion

To understand the evolutionary forces and ecological pressures that shape endophyte assemblages, the delineation of patterns in endophyte biodiversity across spatial scales and the host phylogeny is required. The enthusiasm among microbial ecologists for endophyte biology paired with the tools we now have at our collective disposal, suggests that description of such patterns is within grasp. We hope that our survey inspires others to fill the gaps in knowledge that we report. To that end, we have made the metadata from each study that we consider available (see the Supplemental Material) in hopes that other researchers mine them for additional insights.

## Acknowledgments

Thanks go to Lyra Beltran for assistance extracting data from publications. This review was inspired by conversations with Betsy Arnold, to whom we offer our thanks. We appreciate comments from Leho Tedersoo and two anonymous reviewers that led to a much improved manuscript. JGH was supported by the National Science Foundation EPSCoR grant 1655726. EAG was supported by a Smithsonian Institution Secretary’s Distinguished Research Fellowship, as well as a Smithsonian Environmental Research Center Postdoc Research Fellowship, the Maryland Native Plant Society, the Washington Biologists Field Club, and The New Mexico Idea Network of Biomedical Research Excellence (NM-INBRE).

## Data availability

All scripts and processed data are available at: https://bitbucket.org/harrisonjg/endophytereview/src/master/

## Supplementary Material

### Meta-analysis methods and results

We attempted to perform a meta-analysis to examine relative richness of endophytes among plant tissue types. For this analysis, we omitted those studies that did not standardize observational effort among tissues by either mass or sample count (i.e., the number of samples from each tissue type). We also only considered studies that provided a table describing the counts of each microbial taxon observed within each sample (e.g., an operational taxonomic unit [OTU] table), because these data were required to calculate diversity and richness indices. Out of the 558 studies that examined multiple tissues, nine met these criteria for fungi. For bacteria, only a single study met these criteria, precluding a formal meta-analysis, thus for this taxon we only performed vote counting. We rarefied each OTU table by the minimum number of observations for a sample within that study and calculated richness and exponentiated Shannon’s diversity for each sample. Calculations were performed using the `vegan` R package v2.5-5(Oksanen et al. 2016). A random effects model was used to estimate differences in richness and diversity between tissue types while accounting for among-study variation. Models were implemented using the `metafor` v2.1-0 (Viechtbauer 2010) R package using a restricted maximum likelihood estimation approach.

Across all hosts considered via meta-analysis, we found no significantly supported differences among tissue types in richness or Shannon’s diversity (Figs. S1 &S2).

**Table S1:** Differences among host tissues in fungal (top panel) and bacterial (bottom panel) endophyte richness in woody plants. Each cell in the table provides the number of times the tissue type on that *row* (the focal tissue) had higher richness than the tissue type in that *column* (the comparison tissue) followed by the number of studies reviewed for each comparison in parentheses. Significance was determined using a binomial sign test. For results from herbaceous plants see Table S2, for results from graminoids see Table S3

|  |  | Comparison tissue (Fungi) |  |  |  |  |  |
| --- | --- | --- | --- | --- | --- | --- | --- |
|  |  | Leaf | Root | Stem | Propagule | Flower | Bark |
| Focal tissue | Leaf > | – | 4 (7) | 10 (43)*** | 3 (4) | 2 (3) | 0 (4) |
|  | Root > | 3 (7) | – | 1 (7) | 0 (2) | 1 (2) | 0 (1) |
|  | Stem > | 33 (43)*** | 6 (7) | – | 3 (3) | 2 (3) | 1 (2) |
|  | Propagule > | 1 (4) | 2 (2) | 0 (3) | – | 0 (1) | 0 (0) |
|  | Flower > | 1 (3) | 1 (2) | 1 (3) | 1 (1) | – | 0 (0) |
|  | Bark > | 4 (4) | 1 (1) | 1 (2) | 0 (0) | 0 (0) | – |
|  |  | Comparison tissue (Bacteria) |  |  |  |  |  |
|  |  | Leaf | Root | Stem |  |  |  |
| Focal tissue | Leaf > | – | 2 (7) | 3 (8) |  |  |  |
|  | Root > | 5 (7) | – | 4 (6) |  |  |  |
|  | Stem > | 5 (8) | 1 (6) | – |  |  |  |

\*\*\*p<0.01, \*\*p<0.05, \*p<0.1

**Table S2:**
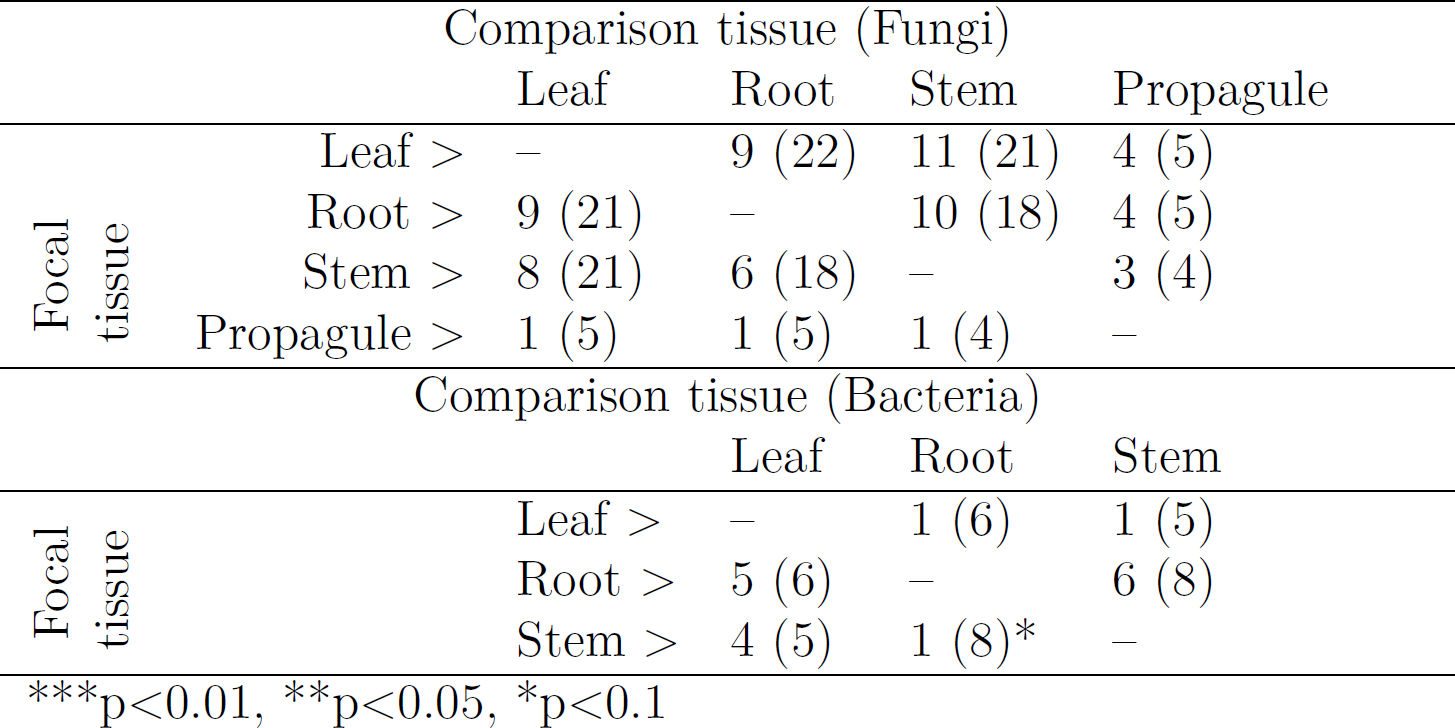
Differences among host tissues in fungal (top panel) and bacterial (bottom panel) endophyte richness in herbaceous plants. Each cell in the table provides the number of times the tissue type on that *row* (the focal tissue) had higher richness than the tissue type in that *column* (the comparison tissue) followed by the number of studies reviewed for each comparison in parentheses. Significance was determined using a binomial sign test. For results from woody plants see Table S1, for results from graminoids see Table S3

|  |  | Comparison tissue (Fungi) |  |  |  |
| --- | --- | --- | --- | --- | --- |
|  |  | Leaf | Root | Stem | Propagule |
| Focal<br>tissue | Leaf > | – | 9 (22) | 11 (21) | 4 (5) |
|  | Root > | 9 (21) | – | 10 (18) | 4 (5) |
|  | Stem > | 8 (21) | 6 (18) | – | 3 (4) |
|  | Propagule > | 1 (5) | 1 (5) | 1 (4) | – |
|  |  | Comparison tissue (Bacteria) |  |  |  |
|  |  | Leaf | Root | Stem |  |
| Focal<br>tissue | Leaf > | – | 1 (6) | 1 (5) |  |
|  | Root > | 5 (6) | – | 6 (8) |  |
|  | Stem > | 4 (5) | 1 (8)* | – |  |

\*\*\*p<0.01, \*\*p<0.05, \*p<0.1

**Table S3:** Differences among host tissues in fungal (top panel) and bacterial (bottom panel) endophyte richness in graminoids. Each cell in the table provides the number of times the tissue type on that *row* (the focal tissue) had higher richness than the tissue type in that *column* (the comparison tissue) followed by the number of studies reviewed for each comparison in parentheses. Significance was determined using a binomial sign test. For results from woody plants see Table S1, for results from forbs see Table S2

|  |  | Comparison tissue (Fungi) |  |  |
| --- | --- | --- | --- | --- |
|  |  | Leaf | Root | Stem |
| Focal tissue | Leaf > | – | 2 (6) | 2 (4) |
|  | Root > | 4 (6) | – | 9 (11)** |
|  | Stem > | 2 (4) | 2 (11)** | – |
|  |  | Comparison tissue (Bacteria) |  |  |
|  |  | Leaf | Root | Stem |
| Focal tissue | Leaf > | – | 1 (6) | 1 (6) |
|  | Root > | 5 (6) | – | 9 (9)*** |
|  | Stem > | 4 (6) | 0 (9)*** | – |

\*\*\*p<0.01, \*\*p<0.05, \*p<0.1

**Figure S1:**
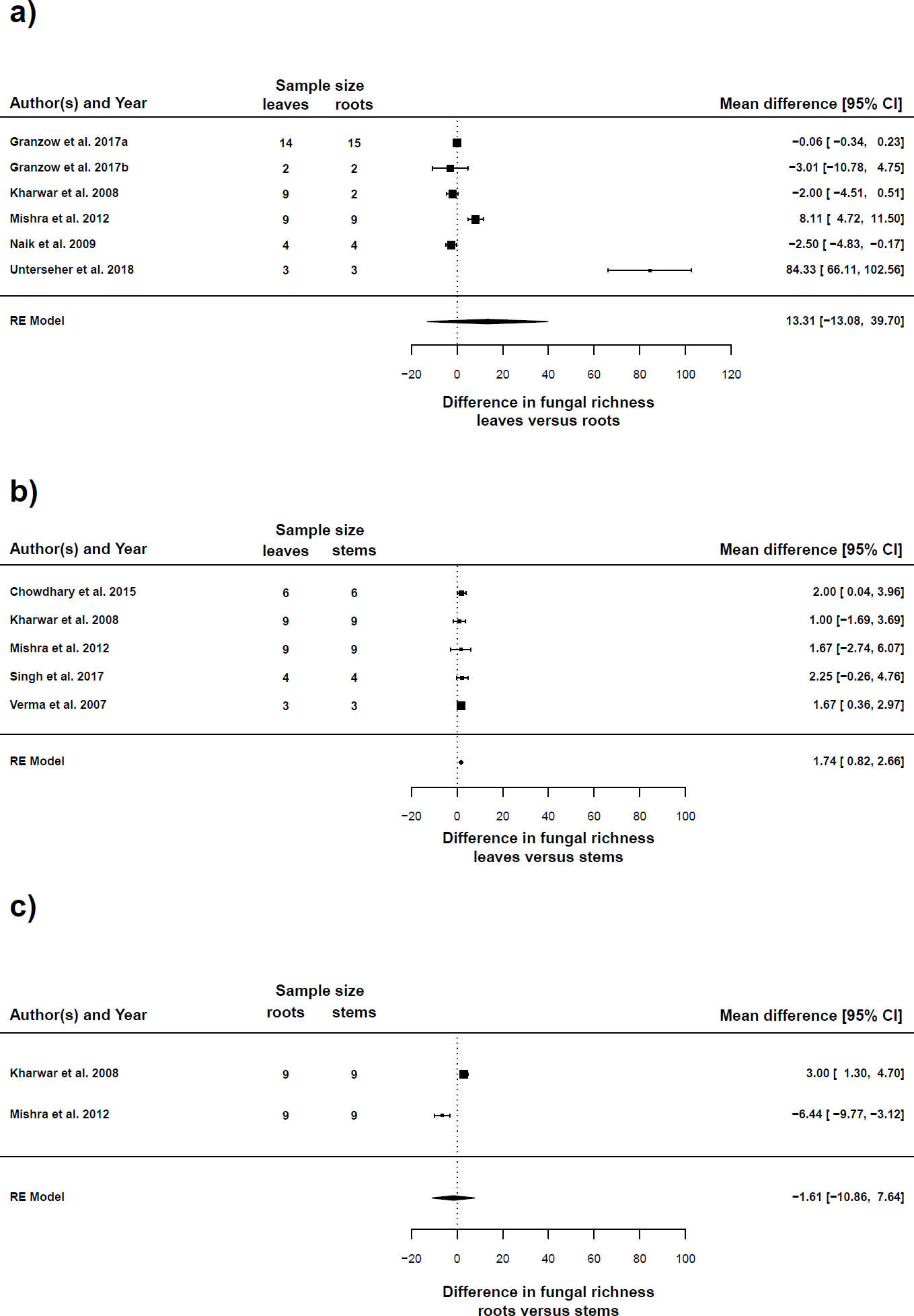
Differences in fungal endophyte richness among host tissues as determined through meta-analysis. Each panel depicts pairwise comparisons between two tissue types. Panel (a) depicts leaves versus roots, panel (b) leaves versus stems, and panel (c) roots versus stems. Mean differences between tissues for each study are shown in the right margins of each plot, with confidence intervals. No model was significantly supported at *p* ≤ 0.05. Results were very similar for Shannon’s diversity and can be seen in Fig. S2. Richness for Unterseher et al. (2018) was higher than the other studies because those authors relied on sequencing data whereas the other studies considered relied on culturing data. Two hosts were studied by Granzow et al. (2017) and results from each host are denoted by letters a and b.

**Figure S2:**
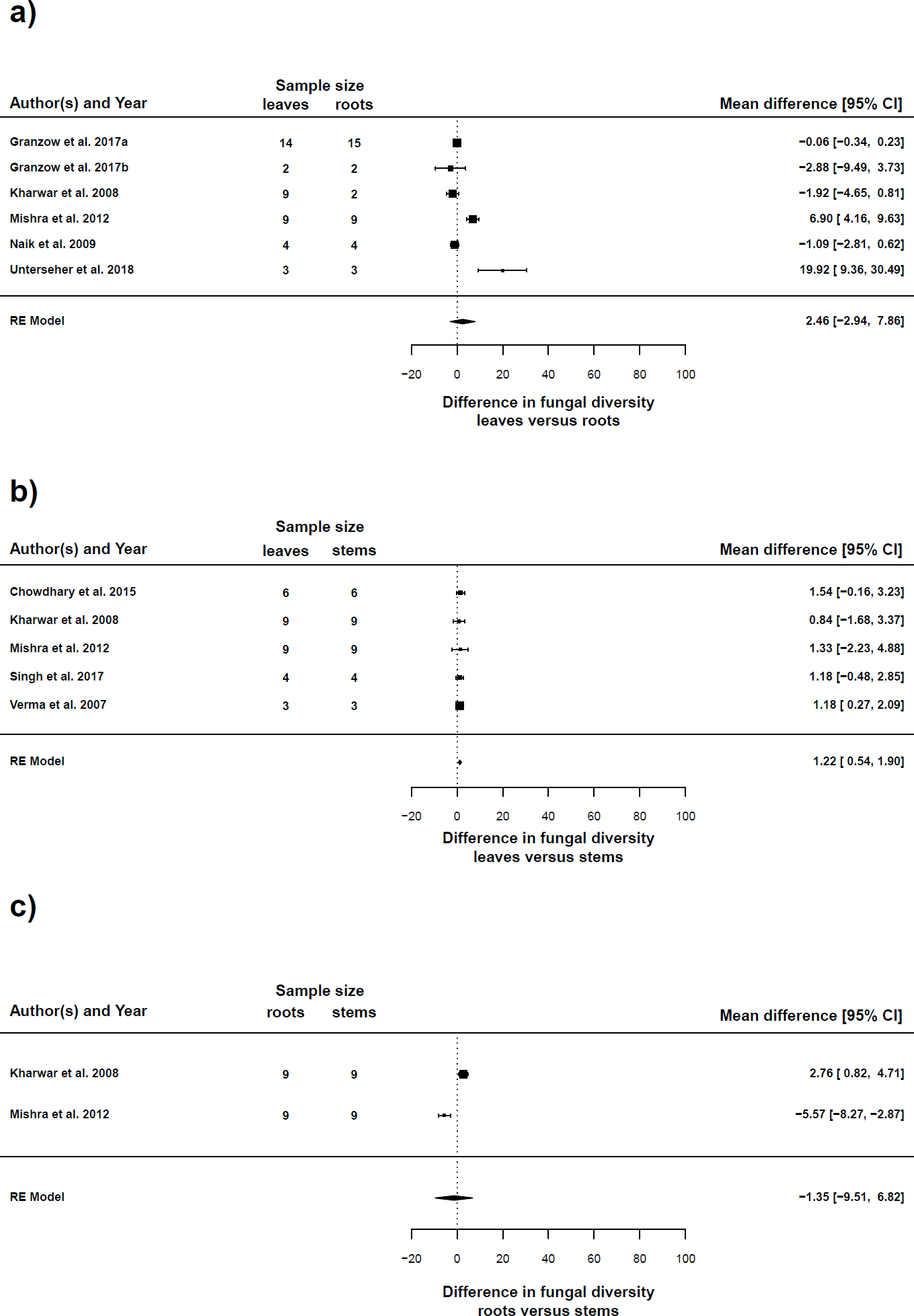
Differences in fungal endophyte diversity (exponentiated Shannon’s entropy) among host tissues as determined through meta-analysis. Each panel depicts pairwise comparisons between two tissue types. Panel (a) depicts leaves versus roots, panel (b) leaves versus stems, and panel (c) roots versus stems. Mean differences between tissues for each study are shown in the right margins of each plot, with confidence intervals. Results were very similar for richness and can be seen in Fig. S1. Diversity for Unterseher et al. (2018) was higher than the other studies because those authors relied on sequencing data whereas the other studies considered relied on culturing data. Two hosts were studied by Granzow et al. (2017) and results from each host are denoted by letters a and b.

